# Results of an attempt to reproduce the STAP phenomenon

**DOI:** 10.1101/028472

**Authors:** Shinichi Aizawa

## Abstract

This reports the results of an attempt by Haruko Obokata to replicate the phenomenon of stimulus-triggered acquisition of pluripotency (STAP), which was first reported in a pair of papers authored by Obokata and colleagues in 2014. The most conclusive evidence for the pluripotency of so-called STAP cells was their purported ability to contribute to chimera formation. In the follow-up trial presented here, putative STAP cells prepared by Obokata were injected into 1154 embryos, of which 671 were recovered. However, the injected cells made no significant contribution in any tissue in any of the embryos developed.

## Introduction

Induced pluripotent stem cells (iPSCs), first reported by Takahashi and Yamanaka using a combination of exogenous genetic factors, have transformed out understanding of the gene regulatory mechanisms underlying cellular pluripotency and differentiation (Takahashi & Yamanaka 2006). This discovery raised the possibility that cellular reprogramming may also be induced by activating endogenous pluripotency genes under certain conditions. In two reports published in *Nature* by Obokata et al (2014a, b), the authors claimed to have observed how “external stimuli such as a transient low-pH stressor reprogram somatic cells into pluripotent cells,” which they referred to as the STAP phenomenon; subsequently, however, after multiple problems were found with the handling and presentation of the data, both papers were retracted.

The present article reports the results of a trial conducted by Haruko Obokata in the RIKEN Center for Developmental Biology (CDB), which was designed to determine whether the STAP phenomenon was in fact reproducible. Obokata was permitted to perform this closely monitored study from July 14 to November 30, 2014 under the supervision of Shinichi Aizawa, head of the Scientific Validity Examination Team, at the direction of the Head Office for Internal Reform organized by the RIKEN President. Unfortunately, I have been unable to contact her since the completion of the trial, or to obtain her agreement to be listed as an author on this article. Nonetheless, given the extraordinary degree of attention and controversy the original STAP publications and research misconduct generated, I feel it is important to report the results of this investigation in the interests of clarifying the scientific record.

The investigation reported here consisted of two types of experiments; preliminary ones conducted without supervision, and formal ones conducted in the presence of expert witnesses. There were no significant differences in the data generated in the preliminary and formal experiments, and all are included together in this report. The experiments were conducted in a new setting, not in the laboratory that Obokata had used for the previous studies described in the retracted *Nature* publications. All reagents, materials, instruments, and experimental spaces were freshly furnished. Obokata was permitted to conduct experiments only in designated rooms, and she did not make any of the analyses herself, other than observations by phase and fluorescence microscopy; she prepared cell aggregates for analysis by other members of the team. In this report, I refer to the studies reported in the papers retracted (Obokata et al., 2014a, b) as “the previous studies” for the sake of brevity. I also refer to the technical tips published by several authors of the original articles for details of the experimental procedure (Obokata et al., 2014c).

## Results

### Frequency of GFP-positive cells from spleen of *Oct-GFP* transgenic mice

Experiments were performed using a transgenic mouse line harboring *GFP* under an *Oct4* promoter (Ohbo et al., 2003); the line is the same as that used in the previous studies (Obokata et al., 2014a, b). The mouse line has been maintained in C57BL/6 background in a homozygous state. Spleens were dissected from homozygous newborn mice (6-8 days old) obtained by crossing a homozygous transgenic female with a homozygous transgenic male, or from hemizygous newborn mice (6-8 days old) obtained by crossing a homozygous transgenic female with a wild type 129 male. Spleen cells were prepared as described previously, but enrichment of CD45-positive cells by FACS sorting was omitted; lymphocytes collected with Lympholyte (Cedarlane Laboratories, Ontario, Canada) were the source of cells used in the experiments.

The stress treatment evaluated was the low-pH condition; no other conditions, such as trituration, were examined. The low-pH conditions included not only the previously reported induction by HCl (Obokata et al., 2014a, b, c), but also that by ATP. Although not described in the previous reports, the ATP treatment had been used most frequently by Obokata et al, and is described in their patent application regarding the STAP process. In brief, the low-pH condition was generated by suspending the 1×10^6^ cells in 494 μl HBSS solution, adding 6 μl 200 mM ATP, and incubating for 15 min at 37 ^o^C in CO_2_ incubator. The low-pH treated-cells were cultured for 6-8 days, and cell aggregates of 50-100 μm showing green fluorescence were identified. Table 1 gives the frequency of such cell aggregates. No apparent difference was found in the frequency of green fluorescent cell aggregates under either of the low-pH conditions (HCl or ATP) or genetic background of mice (C57BL/6 or F1 between C57BL/6 and 129). The observed frequency was approximately 10 green fluorescent cell aggregates per 10^6^ cells seeded; this was approximately 10-fold lower than that in the previous studies. Most green fluorescent cell aggregates also exhibited higher or lower degrees of red fluorescence (Fig. 1). No quantitative determination was made, but about one in three cell aggregates exhibited green fluorescence more intense than red fluorescence. Green fluorescent cell aggregates that exhibited no significant red fluorescence were rare.

**Table 1.** Frequency of green-fluorescent cell aggregates from *Oct*-*GFP* transgenic spleen after low pH treatment

| Mouse background | Treatment | Exp. No. | No. cell aggregates/10<sup>6</sup> cells seeded |  |  |
| --- | --- | --- | --- | --- | --- |
|  |  |  | Average | Minimum | Maximum |
| C57BL/6 | No treat. | 9 | 0 | 0 | 0 |
|  | ATP | 14 | 16 | 4 | 52 |
|  | HCl | 11 | 10 | 3 | 23 |
| F1 (C57BL/6x129) | No treat. | 9 | 0 | 0 | 0 |
|  | ATP | 13 | 12 | 1 | 30 |
|  | HCl | 10 | 10 | 0 | 42 |

**Fig. 1.**
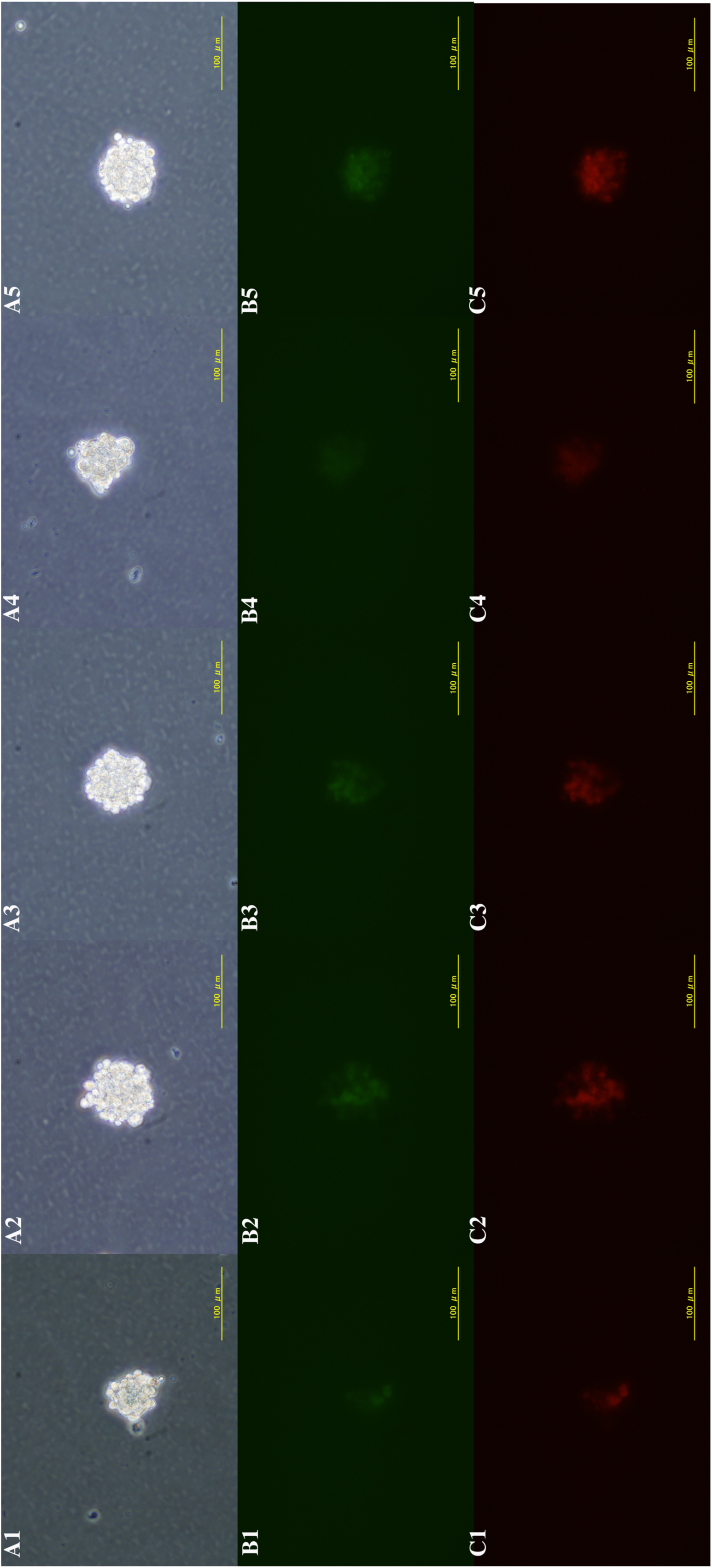
Examples of cell aggregates generated from *Oct-GFP* transgenic spleen by low pH treatment. (A) Phase contrast views of typical five cell aggregates, (B) their green fluorescence and (C) their red fluorescence.

### Chimeric potency of ‘STAP’ cell aggregate

Chimera production was performed with spleens of a transgenic mouse line harboring *GFP* under a *CAG* promoter (Okabe et al., 1997) which were also maintained homozygously in C57BL/6 background; this line is different from the one previously used (Obokata et al., 2014a, b). Cell aggregates of 50-100 μm were selected by their cluster morphology by Obokata and subjected to the chimeric assay. Chimeras were made by members of the Laboratory for Animal Resources and Genetic Engineering, CDB, with expertise in chimera production with ES cells (present affiliation: Animal Resource Development Unit, Biosystem Dynamics Group, Division of Bio-Function Dynamics Imaging, Center for Life Science Technologies (CLST)). The previous report indicated that the generation of chimeras using STAP cells involved a distinct technical approach (Obokata et al., 2014a). “Single cell dispersion by trypsinization, as it is done in the chimera production with ES cells, caused low chimaerism. STAP spherical colonies were cut into small pieces using a microknife under the microscope. Small clusters of the cells are then injected into blastocysts.” In the present study, either an entire cell aggregate or one cut into small pieces by glass capillary, laser beam (XY Clone: Nikko Hansen & Co., Osaka, Japan) or microsurgical knife (K-5310: FEATHER Safety Razor Co., Osaka, Japan) was injected into a host embryo, either E2.5 8-cell stage or E3.5 blastocyst stage embryos of random-bred ICR (Charles River, Tokyo, Japan), as is typical for ES cells. Injected embryos were transplanted into the uterus of pseudopregnant females of the ICR strain, and recovered mainly at E9.5 to judge the contribution of injected cells in each tissue by GFP-green fluorescence (Table 2). Note that in the previous study “small clusters of ‘STAP’ cells are injected into E4.5 blastocysts, and the next day, the chimeric blastocysts were transferred into pseudopregnant females (Obokata et al., 2014a).” In total, 1,154 embryos injected with a cell aggregate were transplanted into a foster uterus, and 671 were recovered. The contribution of injected cells was judged by either GFP green fluorescence in embryos or eye and coat color in live-born mice. No significant contribution of the injected cells was observed in any of these 671 animals. Pluripotency was not examined by injecting putative STAP cells into tetraploid embryos.

**Table 2.**
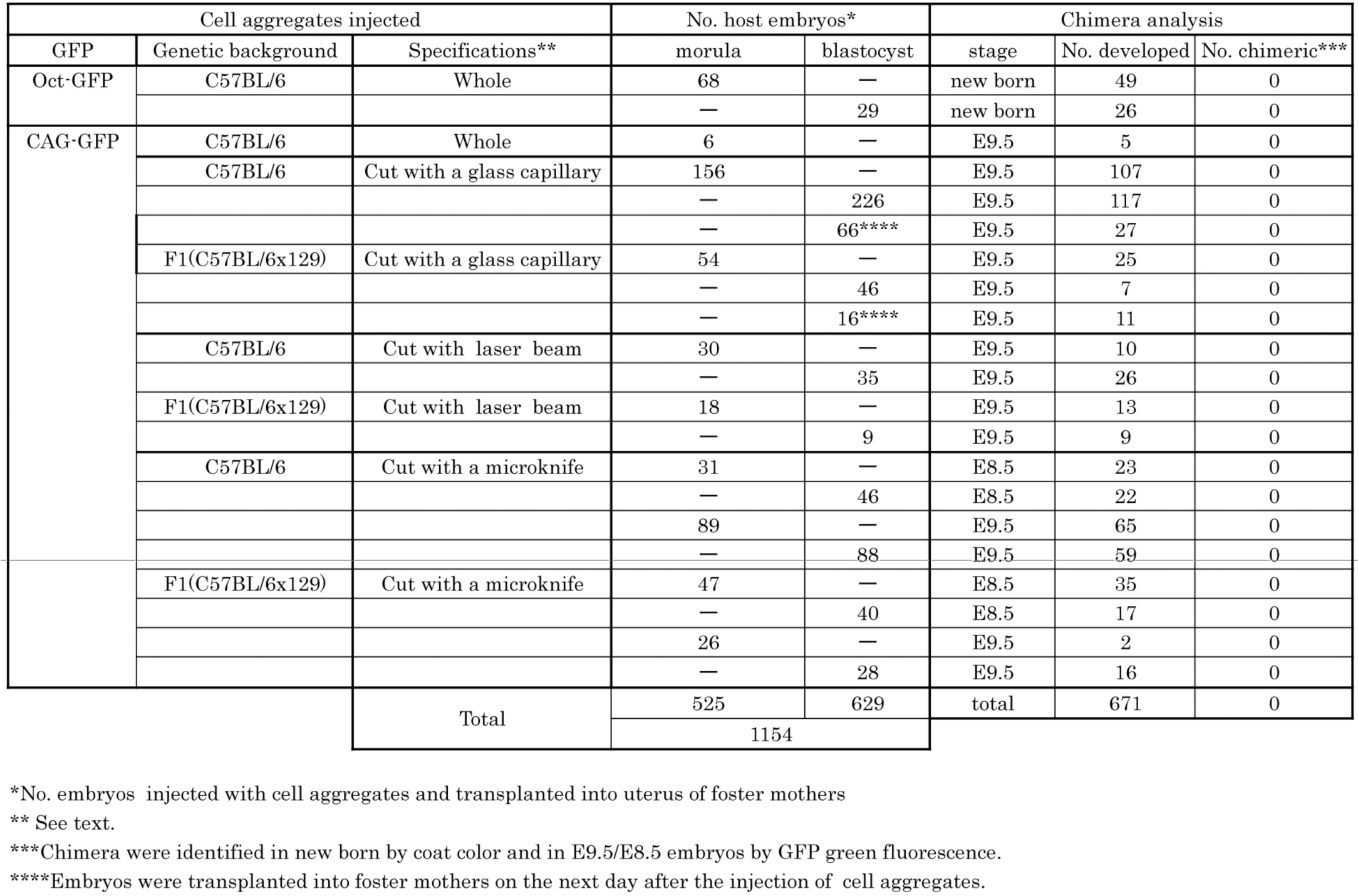
Chimera analysis of pluripotency

## Discussion

One of the central claims in the original reports was that the purported STAP cells had the ability to differentiate into multiple lineages, including germ cells, when placed in a normal developmental environment. However, the present study did not confirm the pluripotency, by chimera production, of cell aggregates prepared by Obokata. In the previous studies, the so-called STAP cells were prepared solely by Haruko Obokata, while the chimera production and the establishment of ES (embryonic stem)-like STAP-SCs and TS (trophectoderm stem)-like FI-SCs were made by Teruhiko Wakayama.

I encourage readers to recognize a number of limitations in the studies reported here, which were conducted under strict time constraints and in the face of considerable, often adversarial, media scrutiny. Unfortunately, it was not possible to have technical advice from Wakayama in the chimera production reported here, and it is unclear whether or to what extent the techniques for chimera production in the present study correspond to those used in the previous studies. Previous studies also examined the pluripotency of purported STAP cells by their potency to generate teratomas in immune-deficient mice. However, more than 10^5^ cells are required to form teratoma subcutaneously in the flank of an immune-deficient mouse using ES or EC (embryo carcinoma) cells, and the process takes about one month. No teratoma formation was examined in the present study, since the frequency of green fluorescent cell aggregates was low and time was limited. Teratoma formation under the kidney capsule, which also takes about two months using blastocyst embryos, was also not examined.

The more critical question is whether and, if so, to what extent the so-called STAP cell aggregates prepared by Obokata in this trial under new experimental conditions recapitulated the STAP cell aggregates reported in the previous study. The frequency of green fluorescent cell aggregates from low pH-treated, *Oct-GFP* transgenic spleen cells was 10-fold less than that in previous studies. The study did not distinguish green fluorescence due to GFP expression from that due to autofluorescence, much less GFP expression resulting from reprogramming from that due to non-specific gene expression in dying cells. The cell aggregates were not characterized in vitro in detail, but the following features were observed:

1. Preliminary FACS analysis of low pH-treated, *Oct-GFP* transgenic spleen cells suggested that the frequency of green fluorescent cells was very low and that the majority of surviving cells were CD45-positive after one week in culture under the conditions used in the present study. In the previous study, CD45 cells were rare and a significant number of green fluorescent cells were observed (Fig. 1c in the previous study (Obokata et al., 2014a)).
2. Preliminary RT-PCR analysis suggested that the majority of the cell aggregates generated in the present study did not express pluripotency markers, in contrast to the report of pluripotency marker expression in the previous study (Fig. 2b in Obokata et al., 2014a), although there were cell aggregates at a low frequency that expressed one or multiple pluripotent markers.
3. Preliminary immunochemical analysis suggested that most of the cell aggregates in the present study did not express pluripotency markers. In contrast to the data shown in Fig. 2a of the previous study, they did not express OCT4, SSEA1, NANOG and E-CADHERIN, (Obokata et al., 2014a).

The possibility cannot be excluded that the experimental conditions used in the present study in some way differed from the previously established optimum conditions for STAP induction. It is my view that it was beyond the scope of this examination to reassign each condition; a definitive answer to the question of whether the previously used conditions for inducing the STAP phenomenon can be indeed established or not must await further study. Nevertheless, I consider it important to report that Haruko Obokata herself failed to reproduce “STAP” cells able to colonize any tissues in a normal developmental environment.

Another reported feature of the STAP phenomenon was that while STAP cells themselves do not proliferate, two types of stem cells could be established from them: STAP-SCs and FI-SCs. However, as Obokata had no experience with stem cell culture, she did not undertake the establishment of these secondary stem cell types in the present study.

## Experimental procedures

Two pieces of spleens from newborn male mice of 6-8 days old were placed in a 15 ml conical tube, minced by scissors into paste, added with 5.5 ml HBSS (GIBCO 14170), mechanically dissociated using a Pasteur pipette and strained through a cell strainer (mesh size 40μm, FALCON 352340) into another conical tube. Five ml of Lympholyte-M (Cedarlane CL5031) was added to the bottom of the tube beneath the cell suspension, and the tube was centrifuged at 1,500g for 20 min. The middle lymphocyte layer was transferred into another tube and centrifuged at 800g for 10 min. The pelleted cells were suspended in 500 μl HBSS, of which 6 μm was subjected to the counting of cell number; in exchange 6 μl 200mM ATP (SIGMA 3377) or diluted HCl (10 μl 35% HCl to 590 μl HBSS) was added to the cell suspension. The cell suspension was incubated at 37^o^C for 15 min in CO_2_ incubator, and then centrifuged at 1,500 rpm for 15min. B27 medium (DMEM/F-12 (GIBCO 11330) supplemented with 1,000 U LIF (ESGRO 1107), 2% B-27 (GIBCO 17504) and 1 μg/ml bFGF (WAKO 060-04543)) was added to the cell pellets to obtain 1×10^6^ cells/ml suspension; one ml of the suspension was plated in each well of a 24 well plate (FALCON 353047) and cultured for seven days to develop cell aggregates.

## Acknowledgements

I would like to acknowledge Ms. Haruko Obokata’s participation and efforts in this study. I am indebted to Dr. Hiroshi Kiyonari and Mr. Kenichi Inoue for chimera production and animal breeding, Laboratory of Animal Resources and Genetic Engineering for animal housing, Dr. Mariko Yamane for RT-PCR analysis and a member of Scientific Validity Examination Team for immunochemical and FACS analyses. I am also grateful to Ms. Kana Bando-Kadowaki, Dr. Go Shioi, Dr. Takaya Abe, Mr. Atsushi Katayama, Mr. Shigekazu Saitou, Mr. Akira Kimura, Mr. Naohiko Oba and Mr. Masahito Hatanaka, for their support to this examination. I deeply thank two senior witnesses outside of RIKEN and seven witnesses from BioResource Center, Center for Integrative Medical Sciences and Brain Science Institute, RIKEN. I thank Mr. Douglas Sipp for critical comments on and copyediting of this report. This examination was supported by the grant for Scientific Validity Examination by RIKEN President.

## References

Obokata, H., Wakayama, T., Sasai, Y., Kojima, K., Vacanti, M. P., Niwa, H., Yamato, M. and Vacanti, C. (2014a) Stimulus-triggered fate conversion of somatic cells into pluripotency. Nature 505, 641–647. Retracted.

Obokata, H., Sasai, Y., Niwa, H., Kadota, M., Andrabi, M., Takata, N., Tokoro, M., Terashita, Y., Yonemura, S., Vacanti, C. A. and Wakayama, T. (2014b) Bidirectional developmental potential in reprogrammed cells with acquired pluripotency. Nature 505, 676–680. Retracted,

Obokata, H., Sasai, Y. and Niwa, H. (2014c) Essential technical tips for STAP cell conversion culture from somatic cells. Nature Protocol Exchange doi:10.1038/protex.2014.008

Ohbo, K., Yoshida, S., Ohmura, M., Ohneda, O., Ogawa, T., Tsuchiya, H., Kuwana, T., Kehler, J., Abe, K., Schöler, H. R. and Suda, T. (2003) Identification and characterization of stem cells in prepubertal spermatogenesis in mice. Dev. Biol. 258, 209–225.

Okabe, M., Masahito Ikawa, M., Kominami, K., Nakanishi, T. and Nishimune, Y. (1997) ‘Green mice’ as a source of ubiquitous green cells. FEBS Letters 407, 313–9.

Takahashi, K. & Yamanaka, S. (2006) Induction of pluripotent stem cells from mouse embryonic and adult fibroblast cultures by defined factors. Cell 126, 663–676.

